# When lysosomes persist: resolving the proton-sponge paradox in nanoparticle-based intracellular delivery

**DOI:** 10.64898/2026.04.29.721565

**Authors:** Indra Van Zundert, Steven Huysecom, Thibo Iven, Sandra Krzyzowska, Vince Goyvaerts, Volker Leen, Johan Hofkens, Hiroshi Uji-i, Beatrice Fortuni, Susana Rocha

**Affiliations:** Molecular Imaging and Photonics, Chemistry Department, KU Leuven, Celestijnenlaan 200F, 3001 Heverlee, Belgium; Chrometra Scientific BV,3470 Kortenaken, Belgium; RIES Hokkaido University, Research Institute for Electronic Science, N20W10, Kita-Ward Sapporo, 0010020, Japan; Institute for Integrated Cell-Material Science (WPI-iCeMS), Kyoto University, Yoshida, Sakyo-ku, Kyoto, 606–8501 Japan

## Abstract

Proton-sponge-active polymers are widely used in nanomedicine to enhance intracellular delivery, yet the mechanism by which they promote cytosolic release of therapeutic cargo remains under debate. Whether these materials drive complete endolysosomal escape or instead alter lysosomal integrity without full nanoparticle release remains unclear. Here we show that polyethylene imine (PEI), a prototypical proton sponge active polymer, induces lysosomal membrane destabilization rather than full nanoparticle escape. Using PEI-coated mesoporous silica nanoparticles as a model delivery system, we show that PEI promotes cytosolic release of small-molecule cargo while nanoparticles remain confined within membrane-enclosed LAMP1-positive compartments. This behaviour arises from the combination of partial lysosomal membrane permeabilization and lysosomal deacidification, which together enable cargo leakage while impairing detection of lysosomes by pH-dependent probes. Our results resolve a long-standing ambiguity in the nanomedicine field and provide a revised mechanistic framework for interpreting endolysosomal escape in intracellular delivery.

**Graphical abstract:** 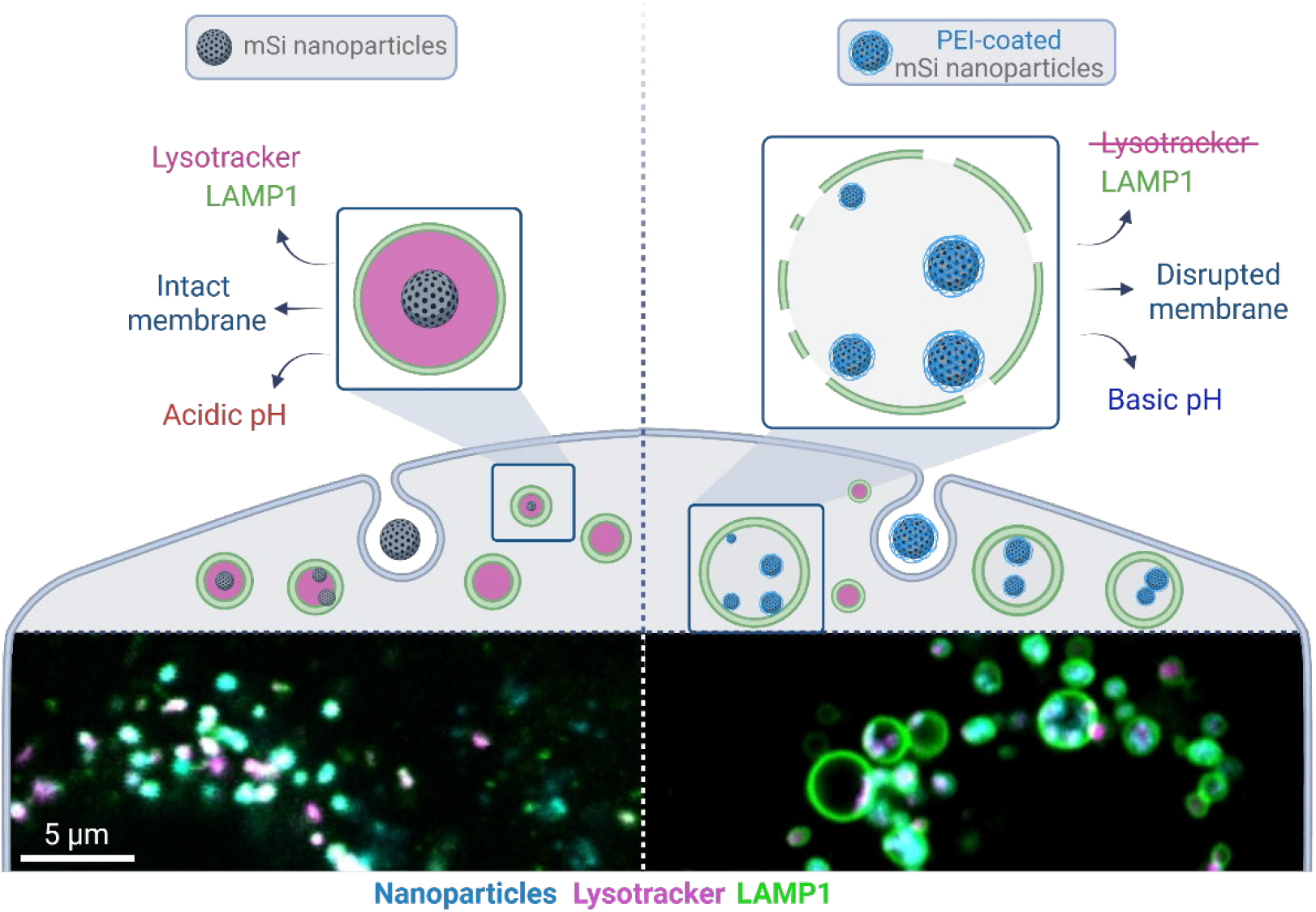

## The unresolved fate of lysosomes in proton-sponge-active delivery

Nanoparticle-based delivery systems are widely used to improve intracellular delivery of therapeutic cargo. Following cellular uptake, however, most nanoparticles remain trapped within endosomes and lysosomes, limiting cytosolic delivery and reducing the activity of many drugs, proteins and nucleic acids.^1–5^ Overcoming this intracellular barrier has therefore become a central objective in nanomedicine development.^6–9^ Cationic, amine-rich polymers such as polyethylene imine (PEI) are frequently incorporated into delivery systems because they enhance apparent cytosolic release and improve downstream biological activity of the delivered cargo.^10–12^ Yet despite their widespread use, the mechanism by which these materials promote intracellular delivery remains under debate.

The leading mechanistic explanation is the proton-sponge effect. In this model, weakly basic groups within the polymer become progressively protonated as endosomal and lysosomal compartments acidify, which is proposed to induce further proton pumping by the vacuolar ATPase. Continued proton influx, accompanied by chloride entry and water uptake, is then thought to cause osmotic swelling^11,13^, membrane destabilization and, ultimately, leakage or rupture of the vesicle. In its strongest interpretation, this process would allow the nanoparticle itself to escape into the cytosol.^14,15^ PEI and related polymers are therefore often described as materials that promote endosomal escape. However, although this hypothesis is pervasive in the literature, direct evidence supporting its key mechanistic steps remains limited and, in some cases, contradictory.^16–19^

Experimental support for proton-sponge-mediated escape has largely relied on indirect functional readouts, including enhanced transfection efficiency of proton-sponge polymer delivered DNA^20,21^ and increased activity of cargo delivered by proton-sponge materials. In parallel, fluorescence colocalization imaging is widely used to infer whether nanoparticles remain trapped within endolysosomal compartments.^22^ Among these approaches, Lysotracker-based imaging is especially common in the field.^23^ In this assay, reduced colocalization between nanoparticles and Lysotracker is often interpreted as evidence of lysosomal escape.^12,24–30^ However, this interpretation relies on two assumptions: first, that lysosomes remain acidic under the conditions being tested, and second, that loss of Lysotracker signal reflects loss of lysosomes rather than loss of acidity. For proton-sponge-active materials, these assumptions are particularly problematic because perturbation of luminal pH is itself central to the proposed mechanism. Under such conditions, reduced Lysotracker signal may reflect impaired acidification rather than loss of the membrane-bound compartment itself.^31–34^ By contrast, lysosomal membrane markers such as LAMP1 report compartment identity independently of luminal pH, underscoring the need to distinguish between loss of acidity and loss of the limiting membrane when assessing endolysosomal escape.

The central unresolved question is therefore not whether proton-sponge-active materials enhance intracellular delivery, but how they remodel endolysosomal compartments during this process. Do these materials induce complete escape of nanoparticles into the cytosol, or do nanoparticles remain membrane-enclosed while endolysosomal integrity and function are selectively perturbed in ways that permit cargo release? Do these compartments remain acidic, or is acidification compromised? Resolving these questions is essential not only for defining the mechanism of proton-sponge-mediated delivery, but also for establishing more rigorous standards to assess endolysosomal escape across nanomedicine.

### Proton-sponge-active nanoparticles promote cargo release

To investigate how proton-sponge-active materials remodel endolysosomal compartments during intracellular delivery, we used mesoporous silica (mSi) nanoparticles coated with PEI as a model system. Mesoporous silica nanoparticles were selected because they represent a widely used carrier platform that can be readily surface-modified and loaded with molecular cargo.^12,35,36^ To enable fluorescence-based tracking in subsequent intracellular imaging experiments, the nanoparticles were synthesized with the fluorescent dye Cy5 incorporated into the silica matrix. Scanning electron microscopy showed that both uncoated and PEI-coated particles were spherical and uniform in morphology Scanning electron microscopy showed that both uncoated and PEI-coated particles were spherical and uniform in morphology (Fig. 1a,b) and had a size of 108.9 and 88.7 nm, respectively (Fig. 1c). Dynamic light scattering (DLS) measurements gave an average hydrodynamic diameter of 165.3 ± 0.87 nm for uncoated particles and 164.2 ± 1.5 nm after PEI coating. The zeta potential shifted from −13.1 ± 0.5 mV for uncoated particles to +51.0 ± 0.5 mV after coating, consistent with successful adsorption of PEI onto the nanoparticle surface. Together, these data establish a well-defined PEI-coated mSi system suitable for probing proton-sponge-associated intracellular effects.

**Figure 1.**
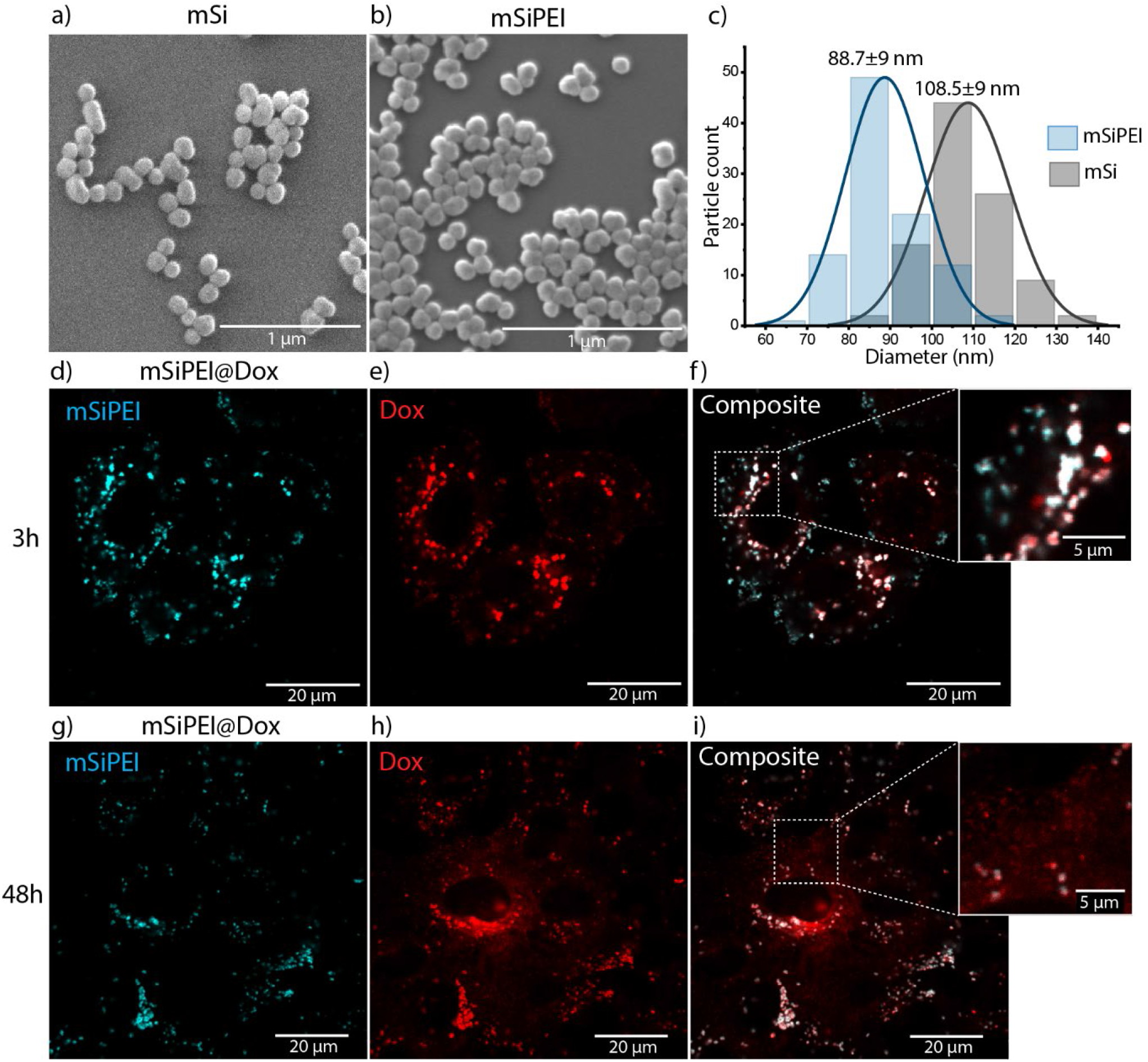
Proton-sponge-active nanoparticles enhance cargo release. **a-c**, Representative scanning electron microscopy images of uncoated Cy5-labelled mesoporous silica nanoparticles (mSi) (**a**) and PEI-coated Cy5-labelled mesoporous silica nanoparticles (mSiPEI) (**b**) and the respective size distribution derived from the measurements (**c**). **d-i**, Representative confocal fluorescence images of cells incubated with doxorubicin (Dox)-loaded PEI-coated nanoparticles and imaged after 3 h (**d-f**) or 48 h (**g-i**). The nanoparticle channel is shown in cyan and doxorubicin fluorescence in red. At 3 h, doxorubicin signal remains predominantly associated with punctate nanoparticle-containing structures (**f**), whereas at 48 h the signal becomes more diffuse, consistent with intracellular cargo release (**i**).

We next assessed whether PEI coating promotes intracellular release of nanoparticle-loaded cargo as previously reported.^12^ To this end, the particles were loaded with doxorubicin (Dox) and incubated with cells, and intracellular drug distribution was monitored over time by confocal fluorescence microscopy using the intrinsic fluorescence of doxorubicin. At the early 3 h time point, doxorubicin fluorescence remained predominantly associated with punctate nanoparticle-containing structures, indicating that most of the cargo was still retained within the carrier shortly after uptake (Fig. 1d-f). By contrast, at 48 h, the doxorubicin signal became markedly more diffuse throughout the cell, consistent with progressive release of the drug from nanoparticle-containing compartments into the cytosolic space (Fig. 1g-i).

Although these observations confirm that PEI facilitates intracellular cargo release, they do not establish whether this results from complete endolysosomal escape or selective perturbation of the compartment. We therefore used this system to investigate how proton-sponge-active nanoparticles alter endolysosomal compartments during cargo release.

### Loss of Lysotracker signal does not indicate nanoparticle lysosomal escape

We next asked whether the apparent cargo release associated with PEI-coated nanoparticles reflects genuine nanoparticle escape from lysosomal compartments. To assess this, we compared two commonly used strategies for identifying lysosomal localization: Lysotracker, an acidotropic dye that accumulates in acidic organelles, and lysosomal-associated membrane protein 1 (LAMP1), a membrane protein enriched in lysosomes and late endolysosomal compartments that reports vesicle identity independently of luminal pH.

Using the conventional Lysotracker-based approach, cells were incubated with uncoated or PEI-coated nanoparticles for 24 h prior to imaging, and nanoparticle overlap with Lysotracker was quantified using Manders colocalization analysis (see Methods). Uncoated nanoparticles showed measureable overlap with Lysotracker (M2 = 0.47), consistent with localization in acidic endolysosomal compartments (Fig. 2a,c). By contrast, PEI-coated nanoparticles showed a significant reduction in overlap with Lysotracker (M2 = 0.24) (Fig. 2b,c). In many studies, such a reduction in Lysotracker colocalization would be interpreted as evidence that the nanoparticles have escaped from lysosomes into the cytosol.

**Figure 2.**
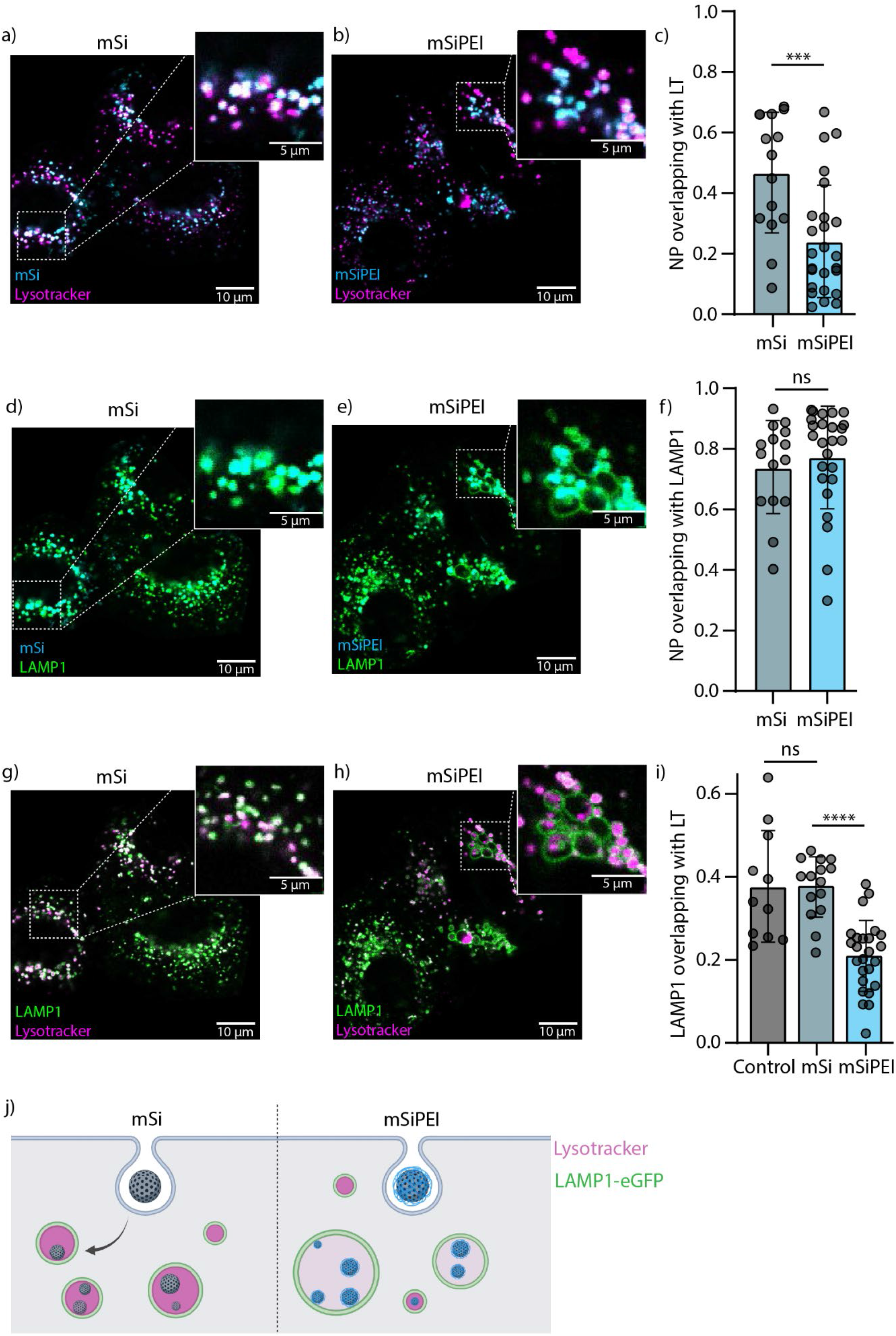
Loss of Lysotracker signal does not indicate nanoparticle escape. **a**,**b**, Representative confocal fluorescence images of cells incubated with uncoated mesoporous silica nanoparticles (**a**, mSi, cyan) or PEI-coated mesoporous silica nanoparticles (**b**, mSiPEI, cyan) and stained with Lysotracker (magenta). **c**, Quantification of nanoparticle colocalization with Lysotracker by Manders analysis, showing reduced overlap for mSiPEI compared with mSi. **d**,**e**, Representative confocal fluorescence images of cells incubated with uncoated nanoparticles (**d**, mSi, cyan) or PEI-coated nanoparticles (**e**, mSiPEI, cyan) in cells expressing LAMP1-eGFP (green). **f**, Quantification of nanoparticle colocalization with LAMP1-defined vesicles by Manders analysis after reconstruction of the enclosed vesicular space from the LAMP1 signal. **g**,**h**, Representative confocal fluorescence images comparing Lysotracker (magenta) and LAMP1-eGFP (green) in untreated cells (**g**) and in cells containing PEI-coated nanoparticles (**h**). **i**, Quantification of the overlap between Lysotracker and LAMP1 by Manders analysis, showing reduced overlap in the presence of mSiPEI. The complementary Manders coefficient is shown in Supplementary Fig. S3. **j**, Schematic summary showing that PEI-coated nanoparticles appear to escape when assessed by Lysotracker, but remain confined within LAMP1-positive lysosomal compartments. Data are shown as mean ± s.d.; *n* > 10 cells per condition. Statistical significance is indicated as ns, not significant; ***P < 0.001; ****P < 0.0001. Scale bars, 10 µm.

We subsequently evaluated nanoparticle localization using fluorescently labelled LAMP1 as a complementary lysosomal marker. Because LAMP1 localizes to the vesicle membrane rather than the lumen, the LAMP1 signal was used to reconstruct the enclosed vesicular space before colocalization analysis; the corresponding workflow is shown in Supplementary Fig. S1. Using this approach, uncoated and PEI-coated nanoparticles showed similarly high overlap with LAMP1-positive vesicles (M2 = 0.74 and 0.77, respectively) (Fig. 2d-f). Thus, although PEI-coated nanoparticles appeared separated from Lysotracker (Fig. 2b), they remained enclosed within membrane-defined LAMP1-positive compartments (Fig. 2e), indicating that reduced Lysotracker colocalization does not necessarily reflect nanoparticle escape.

To directly examine the discrepancy between these two readouts, we next compared Lysotracker and LAMP1 within the same cells. In untreated cells (Supplementary Fig. S2) and in cells incubated with uncoated nanoparticles (Fig. 2g), the fraction of LAMP1-positive vesicles containing Lysotracker was similar (M1 = 0.377 and 0.375, respectively), whereas the fraction of Lysotracker signal overlapping with LAMP1 remained high (M2 = 0.90 and 0.91; Supplementary Fig. S3). This asymmetry reflects the fact that LAMP1 labels a broader population of late endolysosomal compartments, including vesicles that are not sufficiently acidic to accumulate Lysotracker, whereas nearly all Lysotracker-positive vesicles are LAMP1-positive. Accordingly, the relatively low M1 values do not indicate poor agreement between the markers, but rather the broader vesicle population detected by LAMP1.

In the presence of PEI-coated nanoparticles, however, overlap between Lysotracker and LAMP1 decreased significantly (Fig. 2h): the fraction of LAMP1-positive vesicles containing Lysotracker dropped to M1 = 0.20 (Fig. 2i), whereas the complementary coefficient remained relatively high (M2 = 0.82; Supplementary Fig. S3).

Together, these results reveal that PEI-coated nanoparticles disrupt the normal relationship between vesicle acidification and endolysosomal compartment identity. Under normal conditions, Lysotracker and LAMP1 provide largely consistent information about acidic lysosomal compartments. In the presence of PEI, however, LAMP1-positive nanoparticle-containing compartments progressively lose Lysotracker staining, even though the limiting membrane remains present (Fig. 2j) These findings indicate that proton-sponge-active nanoparticles alter endolysosomal physiology in a way that renders Lysotracker an unreliable indicator of nanoparticle escape, and raise two linked mechanistic questions: why do these LAMP1-positive compartments lose Lysotracker staining, and how can cargo be released while nanoparticles remain enclosed?

### Lysosomes remain membrane-enclosed but become partially permeabilized

The discrepancy between Lysotracker and LAMP1 (Fig. 2), together with the observation that cargo is released (Fig. 1) while nanoparticles remain lysosome-confined, suggested that PEI-coated nanoparticles alter lysosomal physiology in a way that permits cargo release without nanoparticle escape. One plausible explanation is that proton-sponge activity induces partial membrane permeabilization: sufficient to allow leakage of small-molecule cargo, but not large enough to release the nanoparticle carrier itself. We therefore asked whether proton-sponge activity induces lysosomal membrane remodeling and permeabilization.

As a first structural readout, we used expansion microscopy to visualize the morphology of nanoparticle-containing, LAMP1-positive compartments at higher effective spatial resolution than conventional confocal microscopy. In this approach, biological samples are embedded in a swellable hydrogel and physically expanded, enabling nanoscale structural features to be resolved using standard fluorescence microscopy.^37^ To retain both the mSiPEI and LAMP1 signals after expansion, LAMP1 was immunolabelled and mSiPEI were conjugated to a gel-linkable label (Ab ExM 405). In untreated cells, LAMP1-positive vesicles appeared as relatively small and well-defined vesicular structures (Fig. 3a,c). By contrast, in cells incubated with PEI-coated nanoparticles, LAMP1-positive compartments appeared enlarged and morphologically irregular (Fig. 3b,d). In particular, nanoparticle-containing vesicles frequently displayed uneven or locally discontinuous LAMP1 signal, consistent with membrane remodelling and localized membrane damage. These observations are in agreement with previously acquired electron microscopy images^12^ and support the idea that proton-sponge-active nanoparticles induce structural stress on endolysosomal membranes and are consistent with swelling and membrane perturbation proposed in proton-sponge-based mechanistic models.

**Figure 3.**
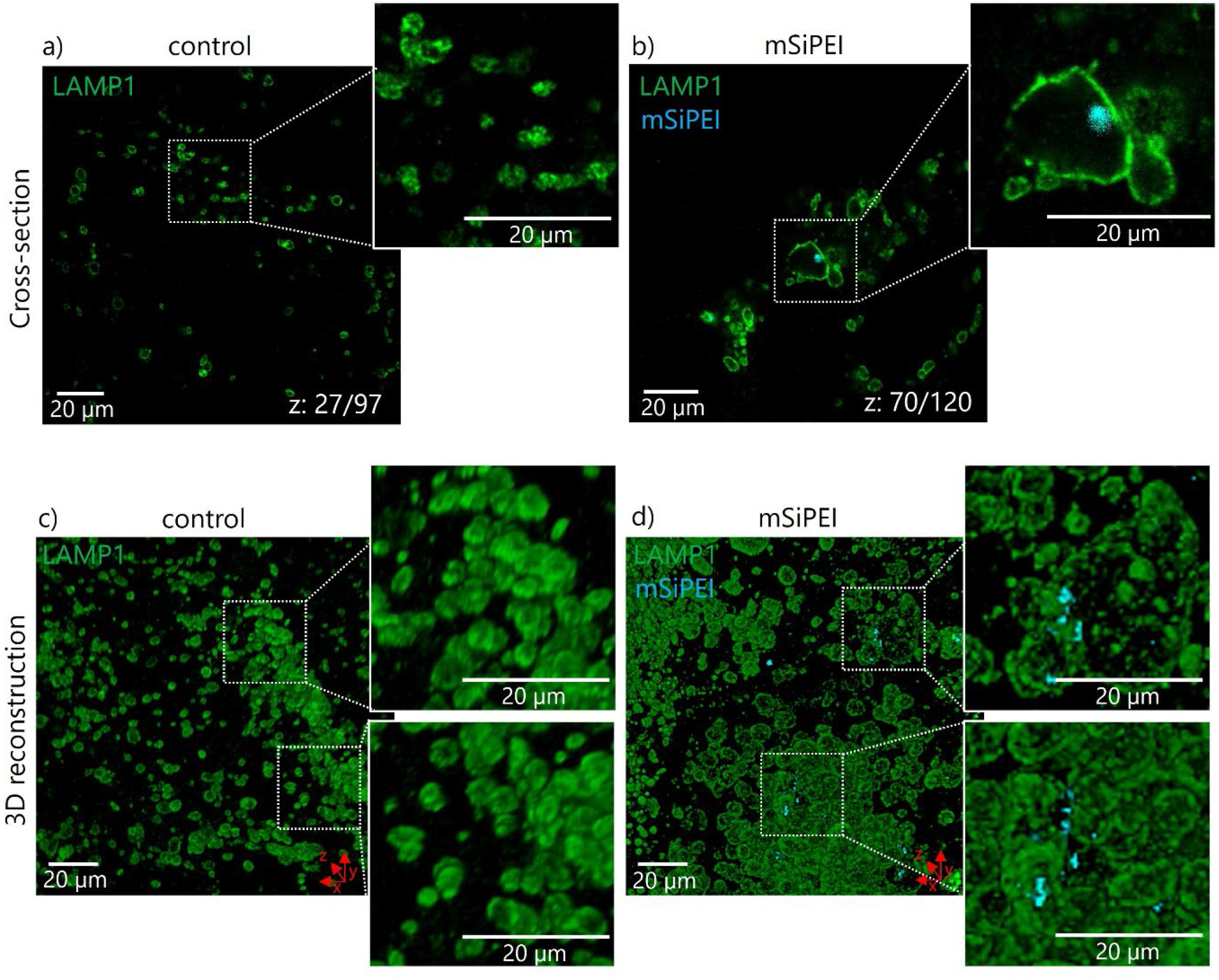
Expansion microscopy reveals altered lysosomal morphology in the presence of PEI-coated nanoparticles. **a**,**b**, Representative single cross-sections of expanded cells expressing LAMP1-eGFP under untreated conditions (plane 27/97) (**a**) or after incubation with PEI-coated mesoporous silica nanoparticles (plane 70/120) (**b**). **c**,**d**, Corresponding 3D reconstructions of LAMP1-positive lysosomes in untreated cells (**c**) and in cells containing PEI-coated nanoparticles (**d**). Compared with untreated cells, lysosomes in mSiPEI-treated cells appear enlarged and morphologically irregular, with locally uneven or discontinuous LAMP1 signal. Nanoparticles are shown in cyan and LAMP1 in green. Scale bars indicate the size after expansion, 20 µm. Using the approximated expansion factor (sevenfold), the scale bars can be corrected to 2.85 µM, representing the size before expansion.

However, expansion microscopy alone does not allow us to conclude unambiguously that these lysosomes are truly permeabilized. If proton-sponge activity causes lysosomal swelling before fixation, LAMP1 proteins may become redistributed over a larger membrane surface, reducing the apparent local signal density. At the increased effective spatial resolution achieved after expansion, this could create the appearance of membrane discontinuities even in the absence of true membrane rupture. We therefore sought an independent readout that directly reports lysosomal membrane damage.

To this end, we performed a Galectin-3 recruitment assay. Galectin-3 is a cytosolic lectin that accumulates on damaged endolysosomal membranes when luminal glycans become exposed to the cytosol (Fig. 4e).^38,39^ Chloroquine, a lysosomotropic weak base that accumulates in acidic lysosomes and perturbs lysosomal pH and membrane integrity, was included as a positive control for lysosomal damage.^40^ In untreated cells and in cells incubated with uncoated nanoparticles, Galectin-3 recruitment to endolysosomal compartments was minimal, indicating little detectable lysosomal damage (Fig. 4a,c,f). By contrast, PEI-coated nanoparticles induced a pronounced increase in Galectin-3-positive vesicles, demonstrating that these particles promote endolysosomal membrane permeabilization (Fig. 4d,f). Chloroquine treatment produced a similar overall response (Fig. 4b,f). Together with the expansion microscopy data, these results show that PEI-coated nanoparticles induce structural remodeling and partial membrane permeabilization of nanoparticle-containing endolysosomal compartments.

**Figure 4.**
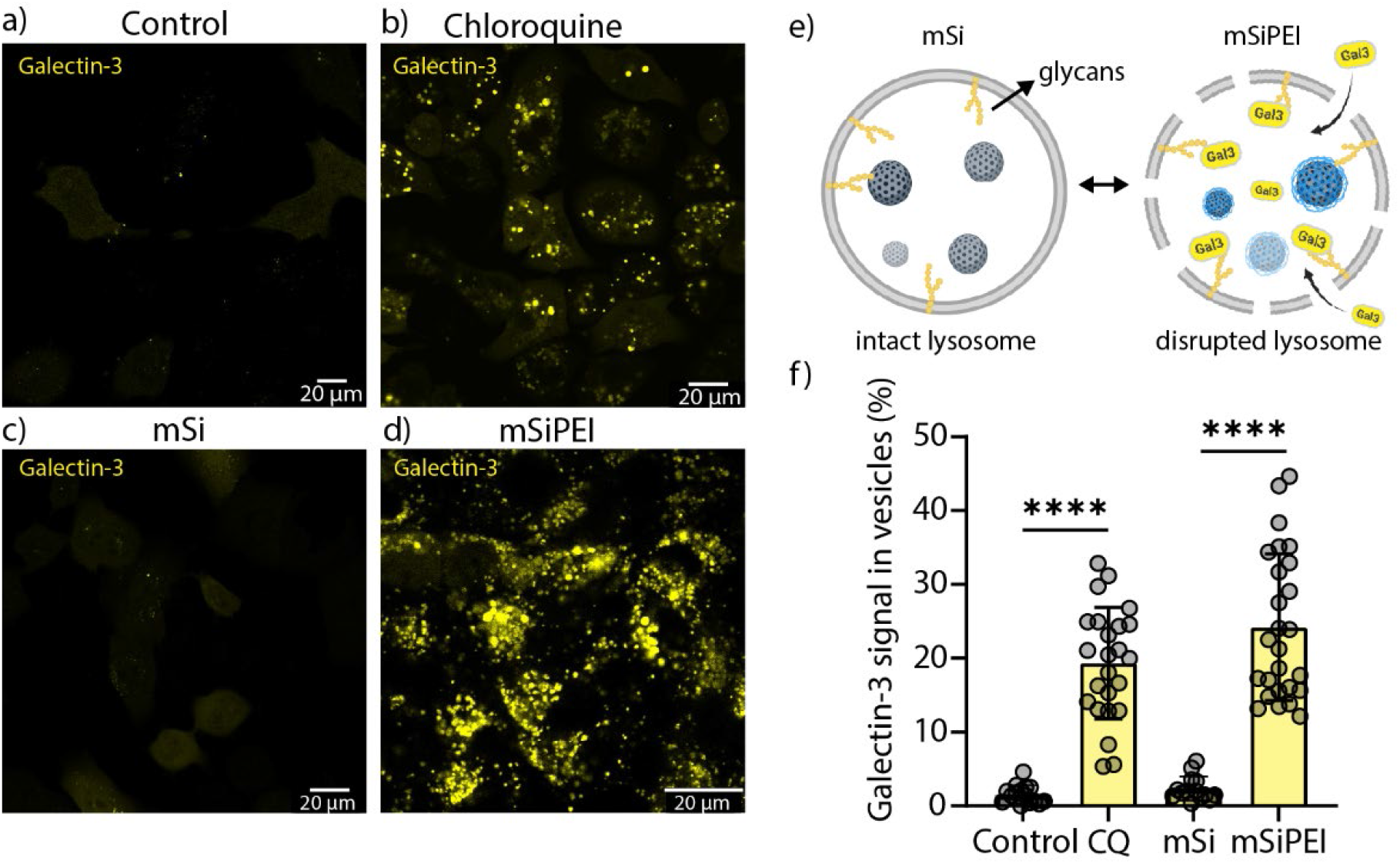
PEI-coated nanoparticles induce lysosomal membrane permeabilization. **a-d**, Representative confocal fluorescence images of A549 cells expressing Galectin-3-mCherry under untreated conditions, additional replicates are shown in Fig. S4 (**a**), after chloroquine treatment (**b**), or after incubation with uncoated mesoporous silica nanoparticles (mSi) (**c**) or PEI-coated mesoporous silica nanoparticles (mSiPEI) (**d**). **e**, Schematic illustration of the Galectin-3 recruitment inside the vesicles. **f**, Quantification of the percentage of Galectin-3 signal recruited to the (broken) lysosomal vesicles in untreated control cells, chloroquine treated cells, or cells incubated with mSi or mSiPEI. Statistical significance is indicated as ****P < 0.0001. Scale bars, 20 µm.

This provides a direct explanation for how small-molecule cargo can access the cytosol while the nanoparticles themselves remain membrane-enclosed. In other words, PEI does not need to drive complete nanoparticle escape to enhance intracellular delivery but perturbation of the lysosomal membrane is sufficient to permit leakage of the cargo.

At the same time, membrane permeabilization alone does not fully explain why LAMP1-positive compartments containing PEI-coated nanoparticles lose Lysotracker staining. Even a partially damaged lysosome would be expected to retain acidotropic dye accumulation as long as luminal acidity is maintained. We therefore next investigated whether proton-sponge-active nanoparticles also alter endolysosomal pH.

### Lysosomal deacidification explains the loss of Lysotracker staining

A central unresolved question in nanomedicine is whether proton-sponge-active materials measurably increase endolysosomal pH. Although these materials are proposed to buffer acidifying compartments, direct evidence for sustained lysosomal deacidification remains limited and contradictory, with many studies reporting little or no detectable pH change.^17^ Determining whether PEI-coated nanoparticles alter lysosomal pH is therefore essential both for evaluating the proton sponge mechanism and for understanding why Lysotracker staining is lost in nanoparticle-containing compartments.

To measure lysosomal pH directly, we used the genetically encoded ratiometric sensor FIRE-pHLy targeted to LAMP1-positive compartments.^41^ FIRE-pHLy consists of a luminal pH-sensitive mTFP1 fused to human LAMP1 and a cytosol-facing, pH-insensitive mCherry reference fluorophore. Because mTFP1 is quenched under acidic conditions, acidic compartments display a low mTFP1/mCherry ratio, whereas alkalinization increases the mTFP1 signal and correspondingly raises the ratio. Thus, the pixel-wise fluorescence ratio provides a direct readout of lumenal pH, with higher ratios indicating reduced acidification (Fig. 5e).

**Figure 5.**
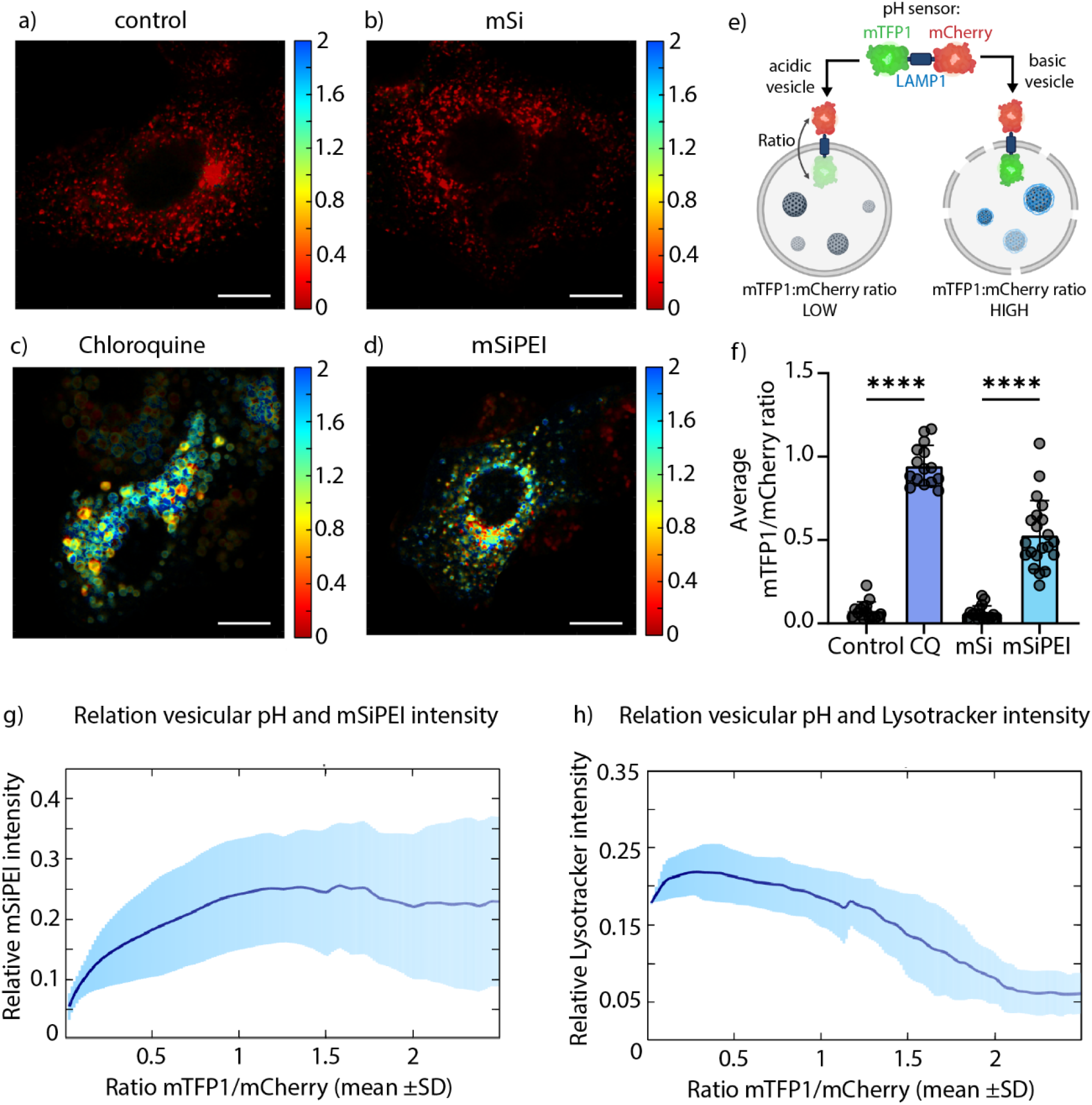
PEI-coated nanoparticles elevate lysosomal pH and impair Lysotracker staining. **a-d**, Representative ratiometric images of cells expressing a lysosome-targeted pH sensor under untreated conditions (**a**), after incubation with uncoated mesoporous silica nanoparticles (**b**, mSi), after chloroquine treatment (**c**), or after incubation with PEI-coated mesoporous silica nanoparticles (**d**, mSiPEI). The color scale represents the average pixel-wise ratio between the luminal pH-sensitive fluorophore (mTFP1) and the reference fluorophore (mCherry). Scale bar, 10 µm. **e**, Schematic illustration of the lysosome-targeted ratiometric pH sensor. **f**, Quantification of the average sensor ratio per cell, showing increased lysosomal pH in chloroquine- and mSiPEI-treated cells compared with untreated and mSi-treated cells. **g**, Pixel-wise analysis of nanoparticle intensity as a function of sensor ratio in mSiPEI-treated cells, showing enrichment of nanoparticle signal in lysosomes with elevated ratio values. **h**, Pixel-wise analysis of Lysotracker intensity as a function of sensor ratio in mSiPEI-treated cells, showing reduced Lysotracker staining at higher ratio values. Data are shown as mean ± s.d. Statistical significance is indicated as ****P < 0.0001.

In untreated cells and in cells incubated with uncoated nanoparticles, LAMP1-positive vesicles displayed uniformly low ratios, consistent with maintained acidic pH (Fig. 5a,b,f). By contrast, chloroquine treatment caused a pronounced increase in the sensor ratio across most vesicles (Fig. 5c,f), consistent with a broad loss of acidification. Cells incubated with PEI-coated nanoparticles also exhibited increased ratios (Fig. 5d,f), demonstrating that proton-sponge-active nanoparticles elevate endolysosomal pH in our system. Quantification of the average sensor ratio per cell confirmed low values for untreated and mSi-treated cells, and significantly higher ratios for chloroquine- and mSiPEI-treated cells (Fig. 5f). Notably, the effect of mSiPEI was more heterogeneous than that of chloroquine: only a subset of vesicles displayed strongly elevated ratios, whereas others remained comparatively acidic. This pattern is consistent with a localized, particle-dependent effect, in which deacidification occurs preferentially in nanoparticle-containing lysosomes. To test this directly, we next correlated compartment pH with nanoparticle intensity. PEI-coated nanoparticle signal was enriched in the high-ratio (deacidified) vesicle population (Fig. 5g), showing that the mSiPEI are preferentially associated with the compartments that have undergone deacidification. This effect was absent when uncoated particles were used (Fig. S5).

Finally, we correlated the pH of LAMP1-positive vesicles to Lysotracker staining intensity (Fig. 5h). In cells treated with PEI-coated nanoparticles, vesicles with elevated FIRE-pHLy ratios (that is, reduced acidification) showed markedly reduced Lysotracker intensity, revealing a clear inverse relationship between compartment pH and Lysotracker staining (Fig. 5h). Thus, the compartments that lose Lysotracker signal remain present and LAMP1-positive, but are no longer sufficiently acidic to retain the dye signal. A similar inverse relationship between sensor ratio and Lysotracker intensity was also observed for chloroquine-treated cells (Supplementary Fig. S6), further supporting the conclusion that reduced Lysotracker staining primarily reflects loss of acidification rather than loss of the membrane-bound compartment itself.

Together, these data show that PEI-coated nanoparticles do not require complete nanoparticle escape to enable intracellular cargo release from lysosomes. Instead, they induce lysosomal deacidification together with partial membrane permeabilization of LAMP1-positive compartments, a combination that permits leakage of small-molecule cargo while abolishing Lysotracker staining. Apparent endosomal escape in proton-sponge-active systems can therefore arise without loss of membrane enclosure.

## Conclusions

Proton-sponge-active materials are widely used in nanomedicine to promote intracellular delivery, yet their mechanism of action has remained unresolved. A central question has been whether these materials drive complete endolysosomal escape of the nanoparticle carrier or instead remodel endolysosomal compartments in ways that permit cargo release without full nanoparticle escape. Here we show that PEI-coated nanoparticles do not undergo complete escape from lysosomes. Instead, they remain membrane-enclosed while inducing partial membrane permeabilization together with reduced acidification.

This mechanistic distinction explains two hallmark observations that are frequently interpreted as evidence of endosomal escape. First, partial membrane permeabilization is sufficient to allow release of small-molecule cargo into the cytosol while retaining the nanoparticle carrier within the compartment. Second, reduced acidification abolishes Lysotracker accumulation, creating the appearance of nanoparticle escape even though LAMP1-positive compartments persist. Our results therefore resolve a long-standing ambiguity in the interpretation of proton-sponge associated delivery.

More broadly, our findings show that apparent endolysosomal escape cannot be inferred from pH-dependent probes alone in systems that perturb lysosomal acidification. Instead, rigorous assessment requires orthogonal evaluation of cargo release, membrane integrity and lysosomal pH. Together, our data support a revised mechanistic framework in which proton-sponge-active materials promote intracellular delivery through partial membrane permeabilization and deacidification, rather than through complete nanoparticle escape. This distinction has important implications for how intracellular delivery systems are designed, evaluated and benchmarked across the nanomedicine field.

## Materials and Methods

### Materials

Low molecular weight polyethylenimine (PEI, Mw ≈1.2 kDa, 50% w/v in H_2_O), Cyanine 5 (Cy5-NHS ester), APTES ((3-Aminopropyl)triethoxysilane), Chloroquine diphosphate powder, Triethanolamine (TEA), 1-octadecene (technical grade), hexadecyltrimethylammonium chloride (CTAC, ≥99%), tetraethyl orthosilicate (TEOS, 98%), hydrochloric acid (HCl, 10 M), Anhydrous ethanol, fetal bovine serum (FBS), Polybrene, N,N’-Methylenebisacrylamide (MBA, ≥99.5%), Ammonium persulfate (APS, ≥98.0%), Anhydrous dimethyl sulfoxide (DMSO), Bovine Serum Albumin (BSA, 7.5% solution in DPBS) were purchased from **Sigma Aldrich**. AB ExM 405, AB ExM 570, and Acrylink NHS (Acryloyl-X, SE) were purchased from **Chrometra Scientific BV**. (Rhodamine 570–conjugated secondary antibodies were prepared in-house.). Anti-eGFP antibody [9F9.F9] (ab1218) and Goat anti-Rabbit IgG H&L antibody (ab6702) were purchased from **Abcam**. Goat anti-Mouse IgG (H+L) Cross-Adsorbed Secondary Antibody (AB_228299), Dulbecco’s modified eagle medium (DMEM), gentamicin, phosphate buffered saline (PBS, no calcium, no magnesium), paraformaldehyde ((v/v 16%, methanol-free), Trypsin-ethylenediaminetetraacetic acid (Trypsin-EDTA, 0.5% solution, no phenol red), Hank’s balanced salt solution (HBSS, no phenol red), Triton™ X-100, UltraPure™ Low melting temperature agarose (1%), GlutaMax™ LysoTracker™ Red, LysoTracker™ Deep Red, DAPI, Acrylamide (AA, 40% solution), Guanidine hydrochloride (99.5%) and Ethylenediaminetetraacetic acid (EDTA, 0.5 M, molecular biology grade) were purchased from **ThermoFisher Scientific**. FuGENE® HD was purchased from Promega Corporation. N,N,N’,N’-Tetramethylethylenediamine (TEMED, >98.0%) was purchased from Tokyo Chemical Industry. Sodium acrylate (SA, 98.0%) was purchased from BLD Pharmatech Ltd. Tris(hydroxymethyl)aminomethane (TRIS, ≥99.0%) was purchased from Carl Roth. Proteinase K (800 U/mL, biology grade) was purchased from New England Biolabs. Atto647N-DBCO was purchased from Atto Tec. The genes for Galectin-3 fused to mCherry, LAMP1 fused to eGFP and the mTFP1/mCherry ratiometric pH sensor (pLJM1-FIRE-pHLy) were obtained from **Addgene** (#85662, #34831 and #170775, respectively) and inserted into a lentiviral vector (#79121). The packaging (pCMV-dR8.2 dvpr, #8455) and VSV-G envelope plasmids (pMD2.G, #12259) for lentivirus production were also purchased via **Addgene**. A549 (adenocarcinomic human alveolar basal epithelial cells) cell line was a gift from Prof. Uji-I (RIES Hokkaido University, Japan) and the Hek 293T cells were a gift from Prof. Mizuno (KU Leuven, Belgium). All chemicals were used without further purification.

### Synthesis of Mesoporous Silica Nanoparticles

Mesoporous silica nanoparticles were synthetized by the biphase stratification method reported by Shen et al.^42^ In short, 0.18 g of TEA was mixed with a solution of 24 mL of CTAC and 36 mL of milli-Q water. This mixture was heated to 60 °C under magnetic stirring for 1 h. Next, 20 mL of TEOS (20 v/v % in octadecene) was slowly added with a syringe and the reaction was kept proceeding overnight. When Cy5 was conjugated to the silica matrix, 4 mg of Cy5-NHS ester was dissolved in 2 mL 99.8% anhydrous ethanol together with 10 µL APTES. This mixture was stirred overnight under inert atmosphere to couple Cy5 to the APTES. After overnight stirring, the solution was added after the addition of the TEOS. The same protocol was followed for the attachment of AB ExM 405 (TFP ester) required for expansion microscopy. After overnight stirring at 60 °C, the reaction was cooled down to room temperature and the nanoparticles were washed with a solution of HCl 1.1 M in water/ethanol (v/v = 1.25:10) with centrifugation-dispersion-sonication cycles to remove CTAC from the pores. Subsequently, the nanoparticles were washed two times with milli-Q water in order to neutralize the pH.

### Doxorubicin loading into mesoporous silica nanoparticles

The pores of the mesoporous silica nanoparticles were loaded with doxorubicin (Dox) for Dox release experiments. Nanoparticles were first dispersed in phosphate buffer (pH 9) to maximize the loading efficiency. To avoid Dox aggregation, the solution containing Dox and mSi was sonicated for 10 min. Next, the solution was stirred for 24 h at 400 rpm. After loading the solution was centrifuged and the supernatant was replaced with milli-Q water and the Dox-loaded nanoparticles were resuspended.

### Nanoparticle Surface Functionalization with PEI

PEI coating was performed by electrostatic adsorption. To coat the nanoparticles with a PEI layer, a 0.75% (w/v) PEI solution (in milli-Q water), adjusted to pH 7 (with 37% HCl) was added to the dye conjugated nanoparticles in a 1:1 ratio in a plastic vial. This mixture was magnetically stirred for 3 h, yielding PEI-coated mSi. After coating, the particles were centrifuged and resuspended in milli-Q water to purify from excess PEI.

### Nanoparticle Characterization

The physicochemical properties of the nanoparticles were characterized using scanning electron microscopy (SEM), Zeta Potential, and Dynamic Light Scattering (DLS). High-resolution SEM images were captured with a field-emission SEM (JEOL JSM-7200F, JEOL Ltd., Japan) microscope operated at 20.0 and 30.0 kV. Samples were sputter-coated with gold prior to imaging. Particle size, size distribution, polydispersity index, and Zeta Potential measurements were performed using a Zetasizer Nano ZS (Malvern Panalytical). All measurements were carried out in milli-Q water at room temperature.

### Cell Culture and Generation of Stable Cell Lines

A549 human lung carcinoma cells were cultured in T25 (25 cm^2^) culture flasks at 37 °C under a 5% CO^2^ atmosphere. Cells were maintained in culture medium (DMEM supplemented with 10% BS, 1% L-glutamax, and 0.1% gentamicin) and passaged using trypsin-EDTA at 80-90% confluency.

Stable A549 cell lines expressing LAMP1-eGFP, Galectin-3–mCherry, both LAMP1-eGFP and Galectin-3–mCherry and mTFP1-LAMP1-mCherry (FIRE-pHLy pH-sensor^41^) were generated using lentiviral transduction. More specifically, 3.3 × 10^6^ Hek 293T cells were seeded in a T25 flask. The next day, the Hek 293T cells were transfected with the packaging (pCMV-dR8.2 dvpr), envelope (pMD2.G) and transfer vector (pHR_SFFV-LAMP1-eGFP or/and pHR_SFFV-Galectin-3-mCherry and pLJM1-FIRE-pHLy) using FuGENE® HD according to the supplier’s protocol. Viruses were produced by the Hek 293T cells for 48 h. After 2 days, the virus-containing medium was aspirated and filtered using a 0.45 µm filter to remove floating Hek 293T cells and cell debris. Next, polybrene was added to the virus-containing medium at a 1:1000 dilution. The filtered virus-containing medium was added directly to the target cells (A549), which were seeded in a 48-well plate at a density of 50 × 10^5^ cells per well the day before. The viruses were incubated with the target cells for 24 h. Transduced cells were expanded and seeded in a 96-well plate at a density of 1 cell per well to obtain monoclonal cultures.

### Nanoparticle Incubation Conditions

Cells were seeded on 29 mm glass-bottom dishes (Cellvis, #1.5 coverslip) and allowed to adhere overnight. Nanoparticles were added to the culture medium at a concentration of 15 µg/mL and incubated with cells for 3 h. Following incubation, nanoparticle-containing medium was removed, cells were washed with phosphate-buffered saline (1x PBS) two times, and fresh medium was added. Cells were then incubated for an additional 24 h (chase period) prior to Lysotracker staining, imaging or fixation (for expansion microscopy). For positive control experiments inducing lysosomal membrane damage, cells were incubated with chloroquine (50 µM) for 24 h.

### Lysotracker Staining

Lysosomes were stained using Lysotracker Red. First, cells were washed three times with HBSS. Next, cells were incubated with Lysotracker at a final concentration of 50 nM (in HBSS) for 30 min at 37 °C prior to imaging. Cells were washed with HBSS before imaging.

### Expansion Microscopy

After nanoparticle incubation, cells stably expressing LAMP1-eGFP were washed twice with 1x PBS, followed by fixation with 4% paraformaldehyde in 1x PBS for 15 min at room temperature. After fixation, the samples were permeabilized and blocked using 0.1% Triton X-100 and 2% BSA in 1x PBS (blocking buffer) for 30 min. Immunostaining was performed by adding mouse an anti-eGFP primary antibody (1:200) diluted in blocking buffer and incubating for 4 h at room temperature. After three PBS washes, the AB ExM 570-conjugated anti-mouse secondary antibody (15 µg/mL) diluted in blocking buffer was added and incubated for 2 h at room temperature. Samples were washed with 1x PBS three times and optionally stained with DAPI (2.5 µg/mL in 1x PBS) by incubating for 5 min at room temperature. Cells were anchored by overnight incubation at room temperature with a 0.1 mg/mL Acrylink NHS solution in 1x PBS (diluted from a 10 mg/mL stock in anhydrous DMSO). Samples were then washed three times with 1x PBS and proceeded to gelation.

TREx gelation solution (10.34% SA, 14.5% AA, 0.005% MBA, 0.15% APS and 0.15% TEMED in 1x PBS) was prepared and maintained on ice to prevent premature gelation.^43,44^ Cells were washed with gelation solution while being cooled on ice. A 30 µL droplet of gelation solution was added to the gelation chamber (microscopy slide covered with parafilm) kept on ice, and the 12 mm coverslip, containing cells, was placed on top of the droplet (cells facing down). After 15 min on ice, the gelation chamber was transferred to an incubator at 37 °C for 1 h. Following gelation, the coverslip containing the gel was carefully removed from the gelation chamber and washed with digestion buffer (50 mM TRIS, 1 mM EDTA, 0.5% v/v Triton X-100 and 800 mM Guanidine hydrochloride in milli-Q water) for 5 min. The gels were then digested overnight at room temperature in 1.5 mL digestion buffer containing Proteinase K (8 U/mL). After digestion, the gels were re-stained with DAPI (2.5 µg/mL in 1x PBS) by incubating for 20 min at room temperature. Subsequently, the gels were transferred to petri dishes and washed with milli-Q water until no further expansion was observed. Upon complete expansion, the gels were cut to the desired shape and size, using a razor blade, and transferred to imaging dishes. To minimize gel drift, samples were surrounded with low melting temperature agarose (1% w/v in milli-Q water).

Expanded samples were pre- and post-stained with DAPI to visualize nuclei. The linear expansion factor was estimated by measuring nuclear diameters before and after expansion and was approximately sevenfold.

### Intracellular pH Measurements

A549 cells stably expressing the FIRE-pHLy pH sensor were incubated with the nanoparticles (mSi and mSiPEI conjugated to Ab ExM 405) and with Chloroquine (50 µM) for 3 h followed by a 24 h chase (no wash after 3 h for the Chloroquine condition). After 24 h, the cells were stained with LysoTracker™ Deep Red as described above. Confocal imaging was performed on live cells as reported in the paragraph “Confocal Fluorescence Microscopy”. Laser powers and acquisition settings for mTFP1 and mCherry were kept constant throughout all the experimental conditions.

### Confocal Fluorescence Microscopy

Confocal fluorescence imaging was performed on a Leica TCS SP8 mini microscope implementing a HC PL APO 63× water immersion objective (NA 1.2) for non-expanded samples and a 25× water immersion objective (NA 0.95) for expanded samples. Distinct diode lasers were used depending on the fluorescent protein or fluorophore. For Cy5 conjugated nanoparticles, 638-nm diode laser was used. For mCherry, LysoTracker™ Red or Rhodamine 6G (after expansion) detection, 552-nm diode laser was used for excitation, while the 488-nm diode laser was used to excite eGFP and Doxorubicin. For AB ExM 405 conjugated particle detection, the 405-nm diode laser was used. Detection was performed with HyD SMD high-sensitivity detectors in standard mode, operating in a detection range of 400 to 800 nm. The detection range was adjusted depending on the fluorophore, with 410-450 nm for AB ExM 405 detection, 500–550 nm for eGFP and mTFP1, 570–600 nm for Doxorubicin, Rhodamine 6G, mCherry and LysoTracker™ Red and 650–750 nm for Cy5 detection. To avoid spectral crosstalk, image acquisition was performed in line (or frame, if required) sequential scanning mode. The images of 1024 × 1024 pixels were acquired at a scanning speed of 200 Hz and line averaging of 3 was applied. For the expanded samples, z-stacks were acquired using a 1 µm step size and a scanning speed of 400 Hz.

### Image Analysis

All image analysis was performed using custom-written scripts in MATLAB (MathWorks). The complete source code is publicly available at https://github.com/StevenHuysecom/SpheroidAnalysis.git.

#### Image Segmentation and Mask Generation

Fluorescence channels were segmented using a custom intensity-based workflow. Images were background-corrected by subtracting a large-radius Gaussian-filtered version of the image to remove low-frequency illumination artifacts, followed by normalization to the maximal pixel value. Manual thresholding was then applied to generate a binary mask. To improve object detection, local intensity maxima were identified and used to refine threshold selection in selected channels. Small objects were removed using size filtering, and connected components were labelled. For vesicular markers (e.g., LAMP1), additional morphological operations, including dilation, hole filling, watershed-based separation, and circularity filtering, were applied to reconstruct closed vesicular structures and define luminal regions. Final masks were converted to binary images and used for downstream quantification.

#### Colocalization and Cross-Correlation Analysis

Colocalization between channels was quantified using both binary mask overlaps. For predefined channel pairs, pixel overlap was calculated and Manders’ colocalization coefficients (M1 and M2) were computed as the percentage of one binary mask contained within the other.

#### Quantification of Galectin-3 Recruitment

Galectin-3 recruitment was quantified by calculating the fraction of total cellular Galectin-3 fluorescence intensity localized within segmented Galectin-3-positive vesicles. Specifically, the sum of raw Galectin-3 intensity values within the corresponding binary mask was divided by the total Galectin-3 intensity per image.

#### Ratiometric pH Analysis and Pixel-Wise Intensity Profiling

Lysosomal pH was estimated using the ratiometric mTFP1/mCherry signal. Background subtraction was performed using morphological opening, followed by median filtering. The pixel-wise ratio (mTFP1/mCherry) was calculated to generate ratio maps. To restrict the analysis to lysosomes containing the sensor, the mCherry channel was segmented using intensity thresholding to identify LAMP1-positive vesicles. The resulting segmentation mask was applied to the ratio maps so that only pixels corresponding to mCherry-positive vesicles were retained. Average ratio values per image were subsequently calculated exclusively from this masked ratio map. For further pixel-wise analysis, only pixels within the masked ratio map were considered. Pixels were binned according to their ratio values, and the median Lysotracker or nanoparticle intensity was computed for each ratio interval. This generated trendlines describing the relationship between lysosomal pH, Lysotracker retention, and nanoparticle localization.

### Statistical Analysis

All quantitative data are presented as mean ± standard deviation. Statistical significance was assessed using a parametric t-test. Statistical significance levels are denoted as ns (not significant), * (p < 0.1), ** (p < 0.01), *** (p < 0.001), and **** (p < 0.0001).

## Acknowledgements

We acknowledge financial support from Research Foundation of Flanders (FWO) research grants (c, G081916N, VS08523N, G0C1821N, and G022724N), postdoctoral fellowships (for BF: 12X1419N and 12X1423N, for IV: 12A6N25N), PhD fellowships (for SH: 11A0S25N, for TI: 1S46825N) and from the KU Leuven (C14/19/079, and C14/22/085).

