## Supplementary information for "When lysosomes persist: resolving the proton-sponge paradox in nanoparticle-based intracellular delivery"

Supplementary info

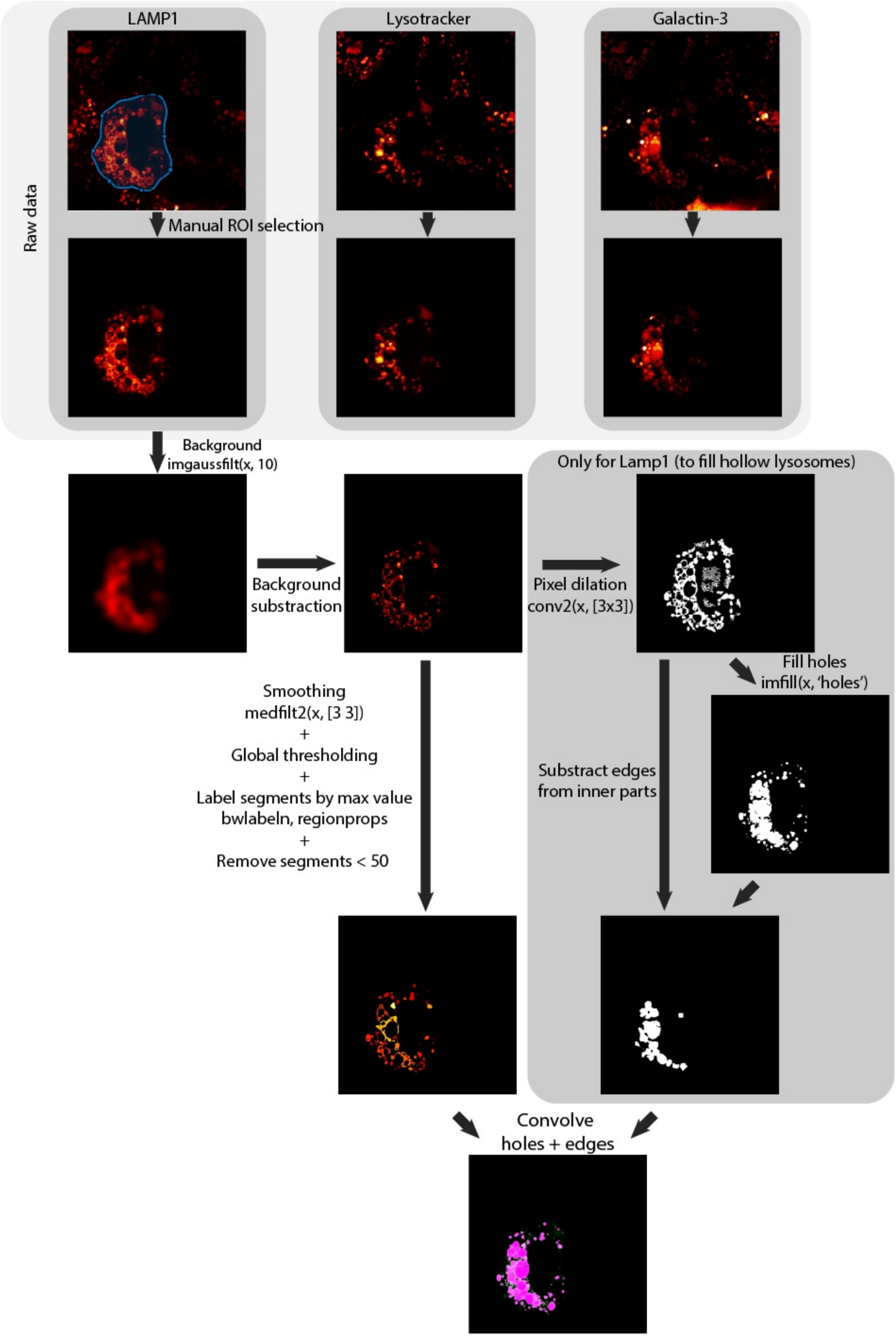


**Figure S1 | Workflow for lysosome segmentation, filling and cross-correlation analysis.** Schematic overview of the image analysis pipeline used for lysosome segmentation and colocalization quantification. Raw fluorescence images of LAMP1, Lysotracker, and Galectin-3 channels are first restricted to manually defined regions of interest (ROIs) corresponding to cells exhibiting detectable signal in all three channels, ensuring analysis of successfully doubletransfected cells only. For each channel, background signal is removed by subtracting a strongly smoothed version of the image (Gaussian filter, σ = 10 pixels), followed by intensity normalization. The background-corrected images are smoothed using a 3 × 3 median filter and globally thresholded to generate initial binary masks of lysosomal structures. Connected components are identified and intensity-normalized by assigning the maximum pixel intensity of each object to all its pixels, and small objects (<50 pixels) are removed. For LAMP1-labeled lysosomes, an additional filling procedure is applied to recover hollow lysosomes. Binary masks are expanded via neighbor-based pixel activation and morphological dilation, followed by hole filling to identify enclosed lumen regions. Interior regions are retained only if they overlap with membrane-derived segments, ensuring that only bona fide lysosomal interiors are included in the final segmentation. For Galectin-3 images, lysosomal segments overlapping with the cell boundary are excluded by detecting and dilating the ROI edge and removing any segmented objects in contact with this boundary. The resulting binary masks represent the final lysosome segmentations for each channel. Pairwise cross-correlation analysis is subsequently performed using these binary masks and the corresponding raw intensity images. Pixel overlap fractions are calculated from the binary masks, Manders' coefficients are computed from masked raw intensity images, and Pearson correlation coefficients are calculated over the union of segmented regions. These metrics provide complementary measures of spatial overlap and intensity correlation between lysosomal markers.

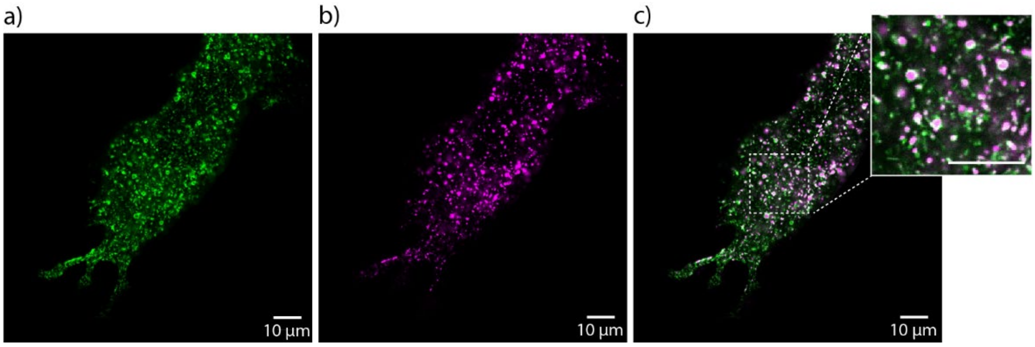


**Figure S2 | Untreated Lysotracker/LAMP1 overlap: a-c**, Representative confocal fluorescence images of A549 cell expressing LAMP1-eGFP (**a**, green) and stained with Lysotracker Red (**b**, magenta) without nanoparticle incubation (**c,** merge) overlap of LAMP1-eGFP and Lysotracker Red.

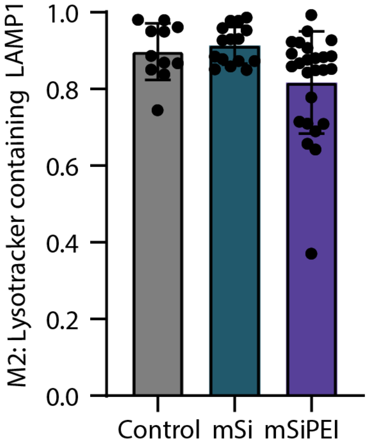


**Figure S3 | Complementary Manders coefficients:** Quantification of the fraction of Lysotracker positive vesicles that are also LAMP1 positive (Manders M2) for untreated control cells, uncoated mesoporous silica nanoparticles (mSi)-treated cells, and PEI-coated mesoporous silica nanoparticles (mSiPEI)-treated cells. Data are presented as mean ± standard deviation.

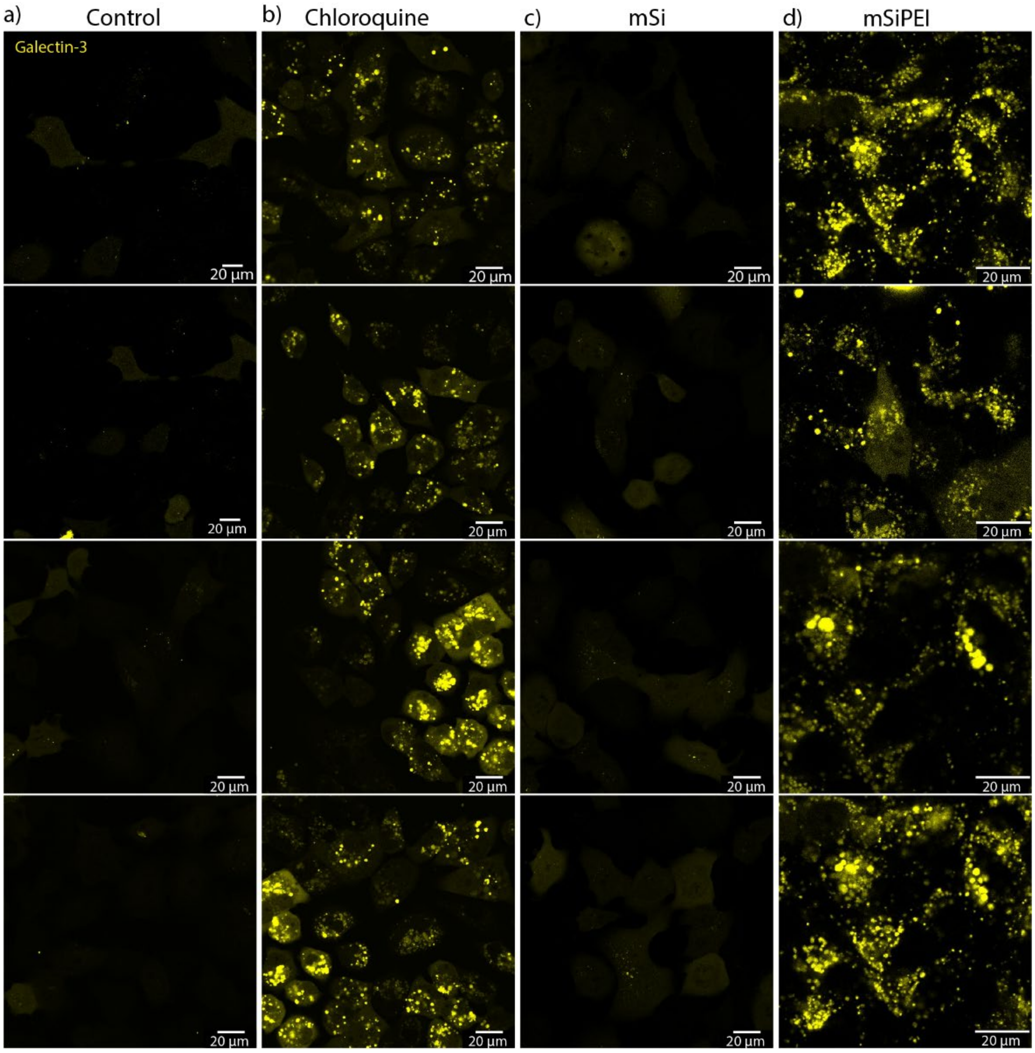


**Figure S4 | PEI-coated nanoparticles induce lysosomal membrane permeabilization. a-d**, Representative confocal fluorescence images of A549 cell expressing Galectin-3-mCherry (yellow) under untreated conditions (**a**), after 50 µM chloroquine treatment (**b**), or after incubation with uncoated mesoporous silica nanoparticles (mSi) (**c**) or PEI-coated mesoporous silica nanoparticles (mSiPEI) (**d**) (4 replicate images shown per condition).

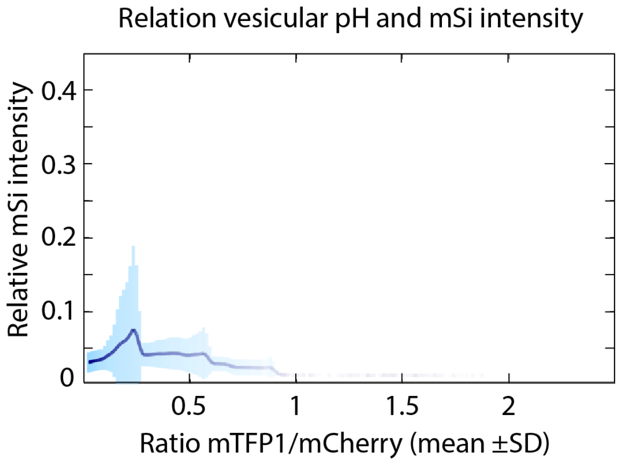

**Figure S5 | Pixel-wise relationships between lysosomal pH and nanoparticle localization.** Pixel-wise analysis of uncoated nanoparticle (mSi) intensity as a function of sensor ratio, showing the relationship between nanoparticle localization and lysosomal pH in the absence of PEI. Transparency indicates the number of pixels contributing to each ratio interval.

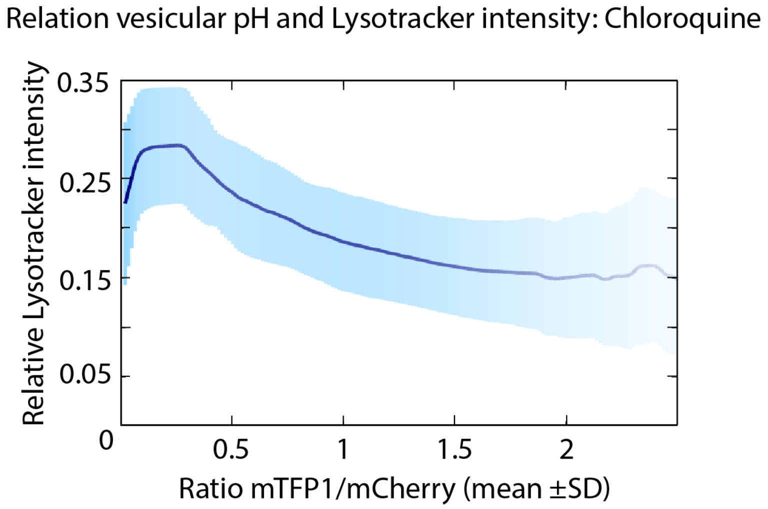


**Figure S6 | Pixel-wise relationships between lysosomal pH and Lysotracker staining.** Pixel-wise analysis of Lysotracker intensity as a function of sensor ratio in chloroquine-treated cells, showing reduced Lysotracker staining at higher ratio values.
